# Simulating PIP_2_-induced gating transitions in Kir6.2 channels

**DOI:** 10.1101/2021.05.20.445012

**Authors:** Michael Bründl, Sarala Pellikan, Anna Stary-Weinzinger

## Abstract

ATP-sensitive potassium (K_ATP_) channels consist of an inwardly rectifying K^+^ channel (Kir6.2) pore, to which four ATP-sensitive sulfonylurea receptor (SUR) domains are attached, thereby coupling K^+^ permeation directly to the metabolic state of the cell. Dysfunction is linked to neonatal diabetes and other diseases. K^+^ flux through these channels is controlled by conformational changes in the helix bundle region, which acts as a physical barrier for K^+^ flux. In addition, the G-loop, located in the cytoplasmic domain, and the selectivity filter might contribute to gating, as suggested by different disease-causing mutations. Gating of Kir channels is regulated by different ligands, like G_βγ_, H^+^, Na^+^, adenosine nucleotides and the signaling lipid phosphatidyl-inositol 4,5-bisphosphate (PIP_2_), which is an essential activator for all eukaryotic Kir family members. Although molecular determinants of PIP_2_ activation of K_ATP_ channels have been investigated in functional studies, structural information of the binding site is still lacking as PIP_2_ could not be resolved in Kir6.2 cryo-EM structures. In this study, we used Molecular Dynamics (MD) simulations to examine the dynamics of residues associated with gating in Kir6.2. By combining this structural information with functional data, we investigated the mechanism underlying Kir6.2 channel regulation by PIP_2_.

## Introduction

Inwardly rectifying K^+^ (Kir) channels are expressed in diverse tissues and regulate physiological processes by setting the cellular resting membrane potential. K^+^ efflux is reduced to different degrees due to block by intracellular Mg^2+^ and polyamines at potentials positive to the K^+^ equilibrium potential (Ho et al. 1993; Kubo et al. 1993; Nichols and Lee 2018). X-ray and cryo-EM structures of several different Kir family members are available, revealing a remarkably conserved pore architecture, despite widely different ligand gating mechanisms. While all Kir channels require phosphatidylinositol-4,5 bisphosphate (PIP_2_) binding for channel activation (D’Avanzo et al. 2010; Hansen, Tao, and MacKinnon 2011; Hilgemann and Ball 1996), gating by many additional ligands is unique. For example, gating of Kir1 and Kir4/5 channels is controlled by pH, Kir3 channels are regulated by Gβγ proteins and Kir6 by ADP/ATP and sulfonylurea receptor subunits (Nichols and Lopatin 1997).

Recent cryo-EM structures of K_ATP_ channels (Ding et al. 2019; Lee, Chen, and MacKinnon 2017; Li et al. 2017; Martin, Kandasamy, et al. 2017; Martin, Yoshioka, et al. 2017; Wu et al. 2018) provide important progress towards understanding the complex gating regulation of this important subfamily, which couples the metabolic state of a cell to its electrical excitability (Hibino et al. 2010; Rorsman and Ashcroft 2018). Structures confirm previous expectations that K_ATP_ channels consist of four Kir6.x pore-forming subunits and four regulatory sulfonylurea receptor (SUR) subunits. Like in Kir2 and Kir3 channels, the pore below the selectivity filter (SF) is lined by two main constrictions: the so-called helix bundle crossing (HBC) gate and the G-loop gate. K^+^ flux through K_ATP_ channels is inhibited by direct interactions of the cytoplasmic Kir6 domain with ATP, while Mg-nucleotide binding to SUR also modulates these channels. Furthermore, channel opening is potentiated by PIP_2_ binding to Kir6, which reduces channel inhibition by ATP (Baukrowitz et al. 1998; Shyng and Nichols 1998). The molecular determinants of PIP_2_ activation in K_ATP_ channels have been investigated in functional studies (Cukras, Jeliazkova, and Nichols 2002; Fan and Makielski 1997; Haider et al. 2007; Männikkö et al. 2011; Pipatpolkai et al. 2020; Schulze et al. 2003; Shyng et al. 2000). However, structural information of the binding site and the gating transitions leading to channel opening are still missing, since all available cryo-EM structures have been solved in the absence of this activator, even though the molecule was present in some cryo-EM experiments (Lee et al. 2017; Wu et al. 2018) (see Table 1 for details). This significantly limits our understanding of the molecular mechanisms by which PIP_2_ and other ligands regulate these channels.

**Table 1:** Available Kir6.2 cryo-EM structures in the PDB compared to a pre-open Kir3.2 x-ray structure

| PDB | State | Experimental Construct | Ligands in Model | Ligands in Experiment | Resolution (Å) | r(L164) (Å) | r(F168) (Å) | r(I296) (Å) | Reference |
| --- | --- | --- | --- | --- | --- | --- | --- | --- | --- |
| 5TWV | Closed | Hamster SUR1, rat Kir6.2 | ATP | glibenclamide, ATP | 6.30 | 1.27 | 1.66 | 2.65 | (Martin, Yoshioka, et al. 2017) |
| 6C3P | Closed (*propeller*) | Human SUR1, human Kir6.2 fusion construct | ATP, ADP, Mg<sup>2+</sup> | ATP, ADP, Mg<sup>2+</sup>, diC8-PIP<sub>2</sub> | 5.60 | 0.97 | 1.24 | 2.36 | (Lee et al. 2017) |
| 6C3O | Closed (*quatrefoil*) | Human SUR1, human Kir6.2 fusion construct | ATP, ADP, K<sup>+</sup>, Mg<sup>2+</sup> | ATP, ADP, Mg<sup>2+</sup>, diC8-PIP<sub>2</sub> | 3.90 | 0.82 | 1.06 | 1.59 | (Lee et al. 2017) |
| 5YW8 | Closed | Hamster SUR1, mouse Kir6.2 fusion construct | ATP<sub>γ</sub>S | ATP<sub>γ</sub>S, glibenclamide | 4.40 | 0.84 | 0.76 | 2.7 | (Wu et al. 2018) |
| 5YW9 | Closed, (T state, *tense*) | Hamster SUR1, mouse Kir6.2 fusion construct | ATP<sub>γ</sub>S | ATP<sub>γ</sub>S | 5.00 | 0.84 | 0.76 | 2.7 | (Wu et al. 2018) |
| 5YWC | Closed | Hamster SUR1, mouse Kir6.2 fusion construct | ADP, Mg<sup>2+</sup> | Mg-ADP, VO<sub>4</sub><sup>3-</sup>, diC8-PIP<sub>2</sub>, NN414 (a KCO) | 4.30 | 0.87 | 0.74 | 2.64 | (Wu et al. 2018) |
| 5YWA | Closed, (R state, *relaxed*) | Hamster SUR1, mouse Kir6.2 fusion construct | ATP<sub>γ</sub>S | ATP<sub>γ</sub>S | 6.10 | 0.89 | 0.71 | 2.91 | (Wu et al. 2018) |
| 5YKF | Closed, (T state, *tense*) | Hamster SUR1, mouse Kir6.2 fusion construct | ATP<sub>γ</sub>S, glibenclamide | ATP<sub>γ</sub>S, glibenclamide | 4.33 | 0.82 | 0.69 | 2.58 | (Wu et al. 2018) |
| 5YWB | Closed | Hamster SUR1, mouse Kir6.2 fusion construct | ADP, Mg<sup>2+</sup> | Mg-ADP, VO<sub>4</sub><sup>3-</sup>, diC8-PIP<sub>2</sub>, NN414 (a KCO) | 5.20 | 0.99 | 0.62 | 2.7 | (Wu et al. 2018) |
| 5YKG | Closed, (R state, *relaxed*) | Hamster SUR1, mouse Kir6.2 fusion construct | ATP<sub>γ</sub>S, glibenclamide | ATP<sub>γ</sub>S, glibenclamide | 4.57 | 0.86 | 0.59 | 2.8 | (Wu et al. 2018) |
| 5YKE | Closed, focus refined TMD + SUR, no CTD | Hamster SUR1, mouse Kir6.2 fusion construct | glibenclamide | ATP<sub>γ</sub>S, glibenclamide | 4.11 | 0.81 | 0.5 | N/A | (Wu et al. 2018) |
| 6BAA | Closed | Hamster SUR1, rat Kir6.2 | glibenclamide, ATP | glibenclamide, ATP | 3.63 | 1.1 | 0.46 | 2.21 | (Martin, Kandasamy, et al. 2017) |
| 6JB1 | Closed, (T state, *tense*) | Hamster SUR1 (Y1209S), mouse Kir6.2 fusion construct | ATP<sub>γ</sub>S, repaglinide, lipids (POPC, PE), digitonin | ATP<sub>γ</sub>S, repaglinide, digitonin, KNtp peptide | 3.30 | 0.96 | 0.38 | 2.41 | (Ding et al. 2019) |
| 5WUA | Closed | Hamster SUR1 (Q608K), mouse Kir6.2 | - | glibenclamide | 5.60 | 1.05 | too narrow | 1.5 | (Li et al. 2017) |
| 3SYA (Kir3.2 ref) | Closed, pre-open | Mouse Kir3.2 crystal structure | diC8-PIP<sub>2</sub>, K<sup>+</sup>, Na<sup>+</sup> | diC8-PIP<sub>2</sub>, Na<sup>+</sup> | 2.98 | 2.95 | 2.45 | 2.96 | (Whorton and MacKinnon 2011) |

Functional studies and modeling strongly support a common structural basis for PIP_2_ regulation in different Kir family members (Cukras et al. 2002; Fan and Makielski 1997; Fürst, Mondou, and D’Avanzo 2014; Haider et al. 2007; Männikkö et al. 2011; Pipatpolkai et al. 2020; Schulze et al. 2003; Shyng et al. 2000). Since PIP_2_ binding sites are well resolved in Kir2 (Hansen et al. 2011; S. J. Lee et al. 2016; Zangerl-Plessl et al. 2019) and Kir3 channels (Niu et al. 2020; Whorton and MacKinnon 2011, 2013), we used this structural information together with functional data to investigate the possible structural basis for K_ATP_ channel regulation by PIP_2_. Specifically, the aim of the present study was to examine PIP_2_-induced gating transitions after unbinding of ATP from the cytoplasmic domain using Molecular Dynamics (MD) simulations. Furthermore, we investigated structural changes of the Permanent Neonatal Diabetes mutation (PNDM) L164P that dramatically alters open state stability of the channel.

## Results and Discussion

### Currently available K_ATP_ structures contain three constriction sites

Fourteen cryo-EM structures were resolved to date, with resolutions ranging from 3.3 to 6.3 Å-resolution. A complete list of currently available Kir6.2 models with included ligands is given in Table 1. To obtain a comprehensive overview of pore dimensions and possible constriction sites, we analyzed pore profiles using the program HOLE (Smart et al. 1996). As shown in Figure 1 and Table 1, three main constriction sites could be identified. Constriction site 1 is located at L164, one helix-turn above the so-called helix bundle crossing (HBC). Constriction site 2, formed by F168 side chains, frames the canonical HBC gate, while I296, a residue associated with the G-loop gate, constitutes the third constriction site. The pore diameter at constriction site 1 ranges from 0.81 to 1.27 Å, in the respective cryo-EM structures. This is in line with previous studies which identified L164 as a narrow, pore-lining site in Kir6.2 (Kurata et al. 2004; Kurata, Marton, and Nichols 2006; Loussouarn et al. 2000, 2001; Proks et al. 2005; Walczewska-Szewc and Nowak 2020). Distances at constriction site 2 range from 0.81 to 1.27 Å, while the pore was slightly wider at I296 (constriction site 3), with distances ranging from 1.5 to 2.9 Å. Overall, none of the structures showed pore radii large enough to enable hydrated K^+^ flux through a continuous pore. Even though recent studies suggest that partially dehydrated K^+^ ions can pass the HBC gate formed by aromatic side chains (Bernsteiner et al. 2019; Black et al. 2020), the constrictions at sites 1 and 3 are formed by hydrophobic residues, rendering K^+^ passage especially at site 1 very unlikely.

**Figure 1:**
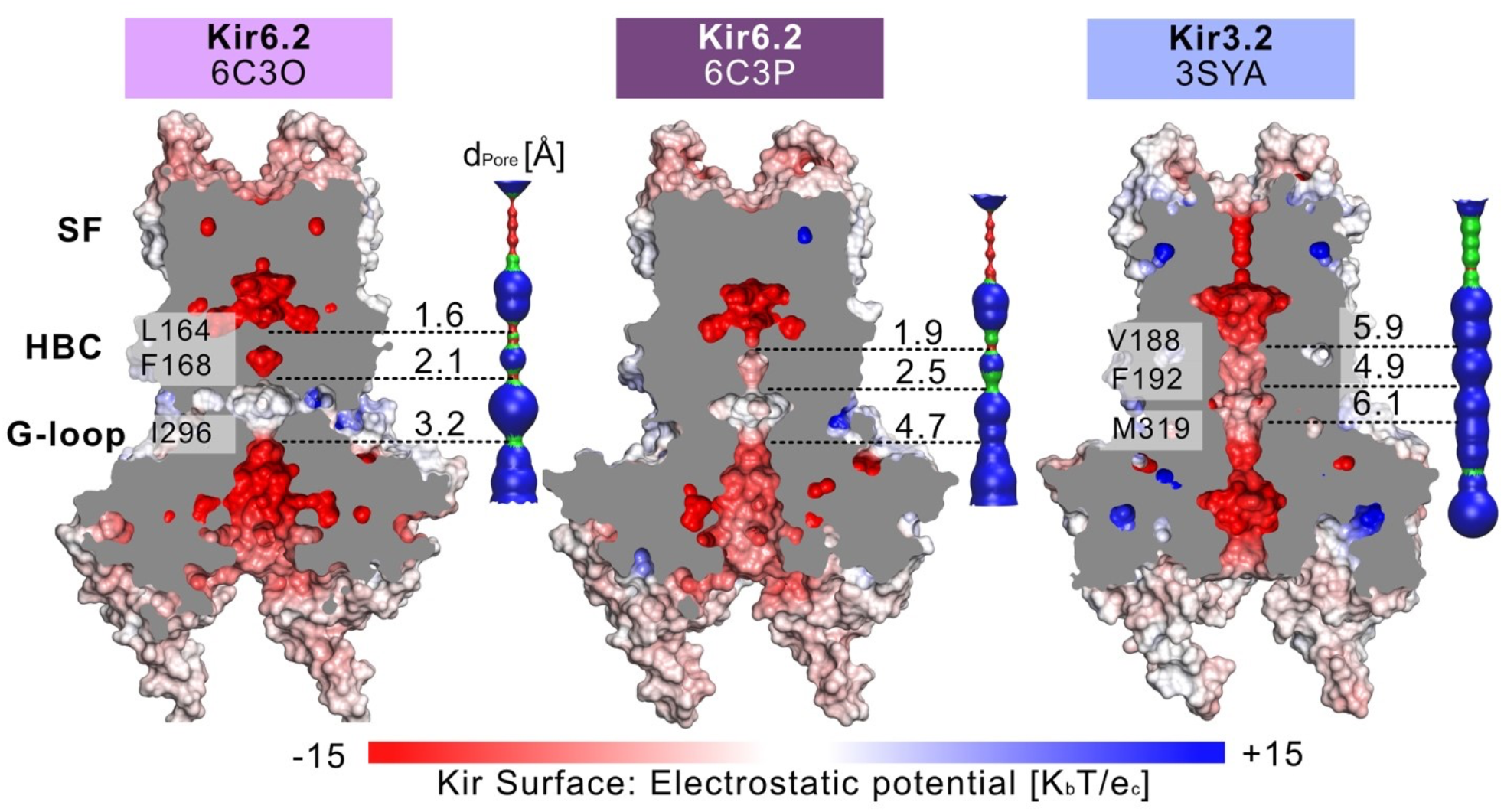
Comparison of the pore dimensions of Kir6.2 structures with Kir3.2. Slices through the pore-forming Kir channel surfaces show three Kir models in the closed state, whereas the pore or Kir3.2 (3SYA) was described as ‘pre-open’. Kir surfaces are colored according to the electrostatic potential. On the right side of each Kir channel, a HOLE profile shows the pore diameter. The three narrowest constriction sites in Kir6.2 below the SF are annotated. In the HOLE profile, red indicates a pore radius too tight for a water molecule. Green suggests space for a single water (0.6 Å < r < 1.15 Å), while blue indicates at least twice the radius of a single water molecule.

### Dynamics of constriction sites in MD simulations

Given our recent success in using MD simulations to provide functional interpretation of conductive state (Bernsteiner et al. 2019)(S. J. Lee et al. 2016)(Zangerl-Plessl et al. 2019) or mutant-induced gating transitions (Linder et al. 2015), we chose to simulate the pore domain of two Kir6.2 structures. Therefore, we selected PDB accession no. 6C3O and 6C3P which were acquired in the presence of the Kir6.2 activators Mg-ADP and PIP_2_ during the experimental setup (Lee et al. 2017) and have quite different radii at the constriction sites as shown in Figure 1. Our recent MD simulations on a Kir3 structure (PDB accession no. 3SYA) revealed that spontaneous wetting of hydrophobic gates could be obtained in classical, atomistic simulations within relatively short time scales (Bernsteiner et al. 2019). Thus, we performed similar MD simulations with the ATP-unbound, closed pore domains of these Kir6.2 structures after placing short-chain PIP_2_ molecules in the respective binding sites, using the 3SYA structure as a template. To differentiate the effect induced by PIP_2_ from inherent protein dynamics, we compared PIP_2_-bound simulations to apo control runs. For simplicity, we will hereinafter refer to these simulations systems as “6C3O” and “6C3P” for the PIP_2_-bound WT simulations, and “6C3O apo” and “6C3P apo” for the control runs without PIP_2_. See table 2 for an overview of simulations.

**Table 2:** Overview of MD simulations

| Name | n | PIP<sub>2</sub> | Length (ns) | Electric Field (mv/nm) |
| --- | --- | --- | --- | --- |
| 6C3O | 5 | + | 1000 | - |
| 6C3O apo | 5 | - | 200 | - |
| 6C3P | 5 | + | 1000 | - |
| 6C3P apo | 5 | - | 200 | - |
| 6C3P | 1 | + | 1000 | - |
| 6C3P | 1 | + | 1000 | 40 (outward) |

To assess the pore’s lumen at the constriction sites, we monitored the minimum distance of each residue between opposing subunits of 5 x 1μs simulations over time, as shown in Figure 2A-C. L164, forming constriction site 1, sampled minimal distances between fully closed and slightly widened apertures, with average minimum distances of 3.4 and 4.4 Å in 6C3O and 6C3P, respectively. Still, both systems remain not only too narrow for hydrated K^+^ ions (6.6 Å (Conway 1981)), but also for water molecules to pass. The pore radii at the canonical HBC gate are wider in both systems. As shown in Figure 2B, a bimodal distribution at F168 causes an asymmetrical pore geometry in both systems. Minimum distances between two opposing subunits frequently sampled 2.6 and 5.8 Å in the 6C3P structure, while the wider pair exhibited distances around 6.7 Å in 6C3O. Despite starting from a narrower HBC gate (1.06 A), pore radii increased to 10 Å in 6C3O, leading to sporadic wetting of the lower HBC gate. However, no persistent wetting in this region could be observed within 1 μs simulation time, as shown in supplementary movies 1 and 2. Thus, PIP_2_ influences the gate diameter of the HBC gate. This is particularly evident for the 6C3O structure, which occasionally permits distances wider than the first K^+^ solvation shell. The G-loop gate diameter increased in both structures, peaking at 9.3 Å in 6C3O and 10.2 Å in 6C3P, which lead to continuous solvation in all simulations (Figure 2C, and supplementary movie 1 and 2). Interestingly, no changes could be observed between PIP_2_-holo and apo simulations at constriction site 1, whereas the bimodal distributions of both systems were shifted towards narrower apertures without bound PIP_2_ in constriction sites 2 and 3.

**Figure 2:**
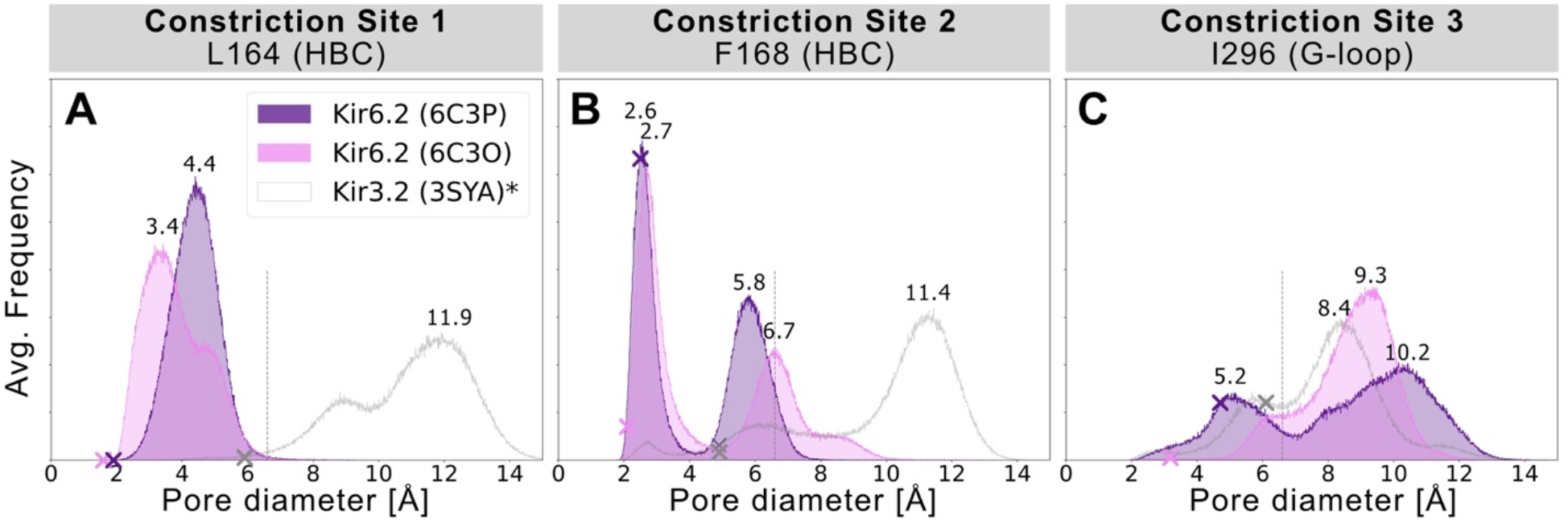
Histograms of pore diameters, sampled in two Kir6.2 systems (6C3O, 6C3P). Minimum distances for three major constriction sites in Kir6.2 were measured in 5 x 1 μs MD simulations for both system between two opposing subunits and averaged over the number of simulations, subsequently. Crosses mark the corresponding distances of the initial cryo-EM state before equilibration and production run, measured with the HOLE program. A vertical line is drawn at 6.6 Å, indicating the time-averaged hydration diameter of K+ (Conway, 1981). Analogous average minimum-distance values, measured in the G-loop gate (narrowest passage in 3SYA), were associated with conduction in previously published Kir3.2 3SYA simulations (Bernsteiner et al., 2019) (*, gray curve).

### Monitoring PIP_2_-induced gating transitions

Given the different behavior of the PIP_2_-bound structures in the simulations, we analyzed the structural changes that led to the short-lived wetting and opening transitions at the HBC gate observed in the 6C3O structure. As previously described in structural and functional studies (Fürst et al. 2014; Hansen et al. 2011; Li et al. 2015; Niu et al. 2020; Poveda et al. 2017), we observed a key structural change in the C-linker, characterized by a PIP_2_-dependent conversion of the loop into a helix. Thus, we monitored changes in the secondary structure of the C-linker over simulation time in both systems, as shown in Figure 3C and 3E, and subplots A, C, E, G of supplementary Figures 2 and 3. Interestingly, there is a ~20% higher helical content (310 helix, corresponding to gray lines in the plots) in the 6C3O C-linker, compared to 6C3P, suggesting a possible connection between the conformation of the C-linker region and the HBC gate. What might cause this early PIP_2_-dependent step towards opening? When comparing residues that render the PIP_2_-binding site in 6C3O and 6C3P, a critical difference becomes apparent. R176 was determined to be a key residue for PIP_2_-dependent activation in Kir6.2 (Baukrowitz et al. 1998; Fan and Makielski 1997; Haider et al. 2007; Pipatpolkai et al. 2020, 2021; Shyng et al. 2000) and other Kir families (Lacin et al. 2017; C. H. Lee et al. 2016; Lopes et al. 2002; Soom et al. 2001; Xie et al. 2005). As shown in Figure 3D and 3F and supplementary Figures 2 and 3, R176 formed stable h-bonds with PIP_2_ in all 5 replicas of the 6C3O system, while h-bonds were missing in one to three subunits of 6C3P simulations. A more detailed analysis suggests a ~ 80% correlation between the R176-PIP_2_ h-bonds and the helical content of the C-linker, corroborating the importance of this interaction for PIP_2_-induced conformational changes. Another important conformational change that has been reported in several previous studies concerns the dynamics of the cytoplasmic domain, including a rotation of the whole domain (Bavro et al. 2012; Clarke et al. 2010; Li et al. 2015, 2017; Linder et al. 2015; Whorton and MacKinnon 2011, 2013; Wu et al. 2018; Zangerl-Plessl et al. 2019). We therefore monitored the rotation of the CTD over simulation time, as shown in supplementary Figure 4. Indeed, there is a stronger relative clockwise rotation observed in the 6C3O simulations (9.2° ± 4.3° std. after 1 μs, n=5, viewed from the extracellular side), compared to 6C3P. Absolute rotation angles of PIP_2_-bound systems converged at ~70° (measured between TMD and CTD of a single protein subunit) in both systems. Nevertheless, fluctuations of the rotation angles render the interpretation of this conformational change rather speculative. Future experiments are necessary to address this question in more detail. Furthermore, an important limitation of our study concerns the lack of the SUR domain, which was not included in our simulation systems, due to poor resolution and many missing residues of these domains. It has to be noted, however, that the N-terminal transmembrane domain of SUR1 modulates PIP_2_ sensitivity in Kir6.2 channels (Pratt et al. 2011; Walczewska-Szewc and Nowak 2020). This might explain why no wetting or widening at the uppermost constriction site was observed in our simulations, rendering the channels essentially non-conductive.

**Figure 3:**
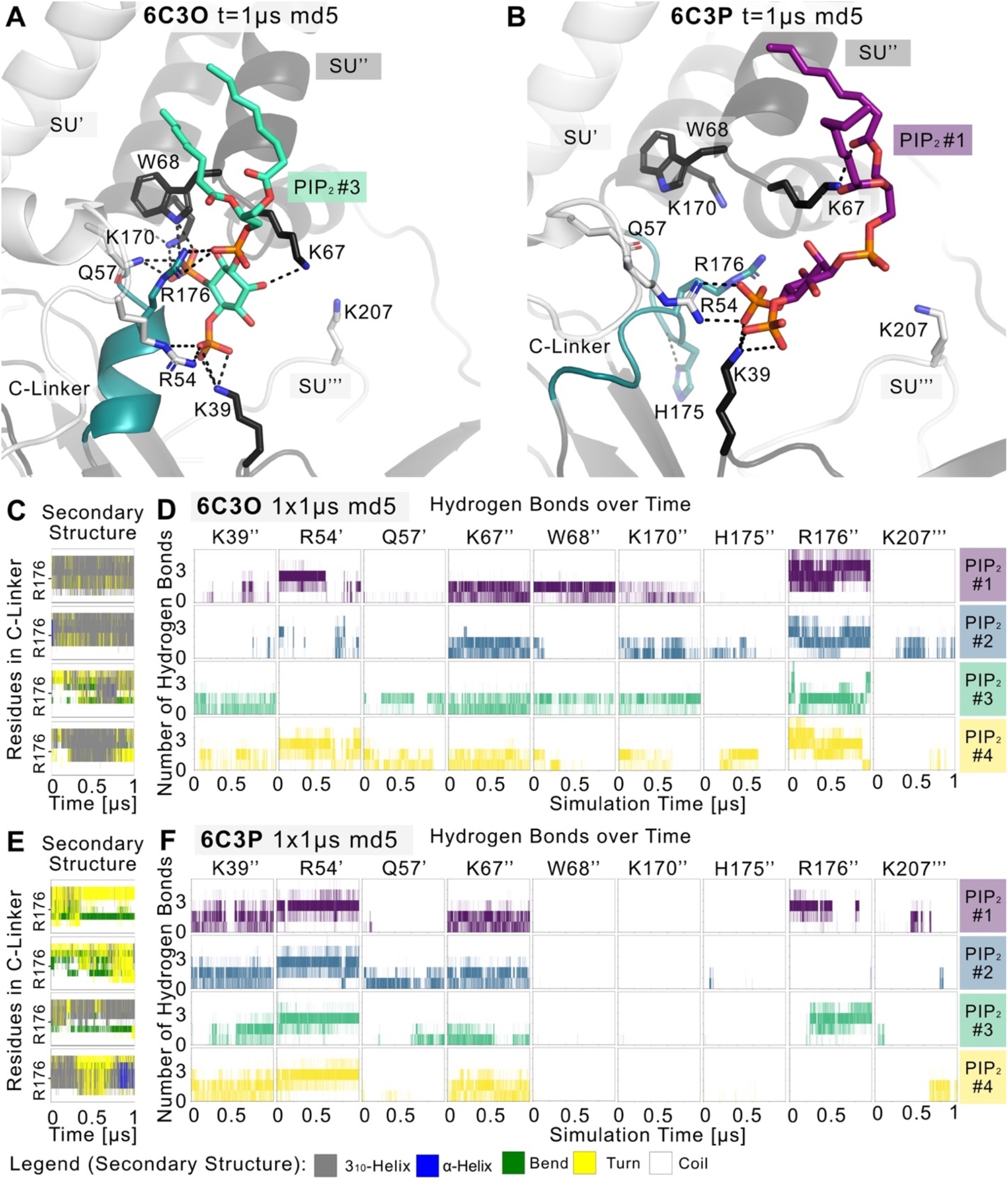
PIP_2_ binding site and PIP2-induced gating changes. Snapshots of polar Kir6.2-PIP_2_ interactions after 1 μs of a representative (A) 6C3O and (B) 6C3P simulation. Relevant residues are shown as sticks, where colors correspond to a protein subunit (SU). The C-linker is colored in teal. (C,E) Plots correspond to four Kir subunits of a representative simulation system. The secondary structure of the (C) 6C3O and (E) 6C3P C-linkers is shown over time, where the y-axis ranges between residues Q173 and T180. The color legend is at the bottom of the figure. (D) Number of hydrogen bonds is shown between PIP_2_ molecules and Kir6.2-residues over time for (D) 6C3O and (F) 6C3P. Subunits in (C,D) and (E,F) correspond.

### Disease mutation L164P strongly influences pore geometry

L164P, a mutation concerning constriction site 1, exhibits very high open-state stability and is insensitive to ATP regulation (Enkvetchakul et al. 2000; Loussouarn et al. 2000, 2001). Heterozygous L164P mutations cause permanent neonatal diabetes by significantly increasing intrinsic open probability (Po) compared to WT channels, while retaining single-channel current amplitudes (Flanagan et al. 2006; Tammaro et al. 2008). To investigate mutation-induced structural changes on the pore we performed MD simulations of the mutant, using the 6C3P system after 1 μs simulation time (Figure 4A). Figure 4B and 4C and supplementary movie 3 show changes in pore solvation during a 1 μs simulation. Due to the lack of the hydrophobic side chain, the minimum distance at constriction site 1 increased to ~10 Å, leading to rapid solvation of the cavity above residue F168 (Figure 4D). Figure 4E shows a shift of the bimodal distribution towards wider average minimum distances (~6.2 Å). Subsequently, constriction site 2 wetted spontaneously with intermittent desolvated periods in both mutant replicas (see supplementary movie 3). Furthermore, the G-loop gate stabilized in an open and fully solvated conformation, peaking at ~10 Å (Figure 4F).

**Figure 4:**
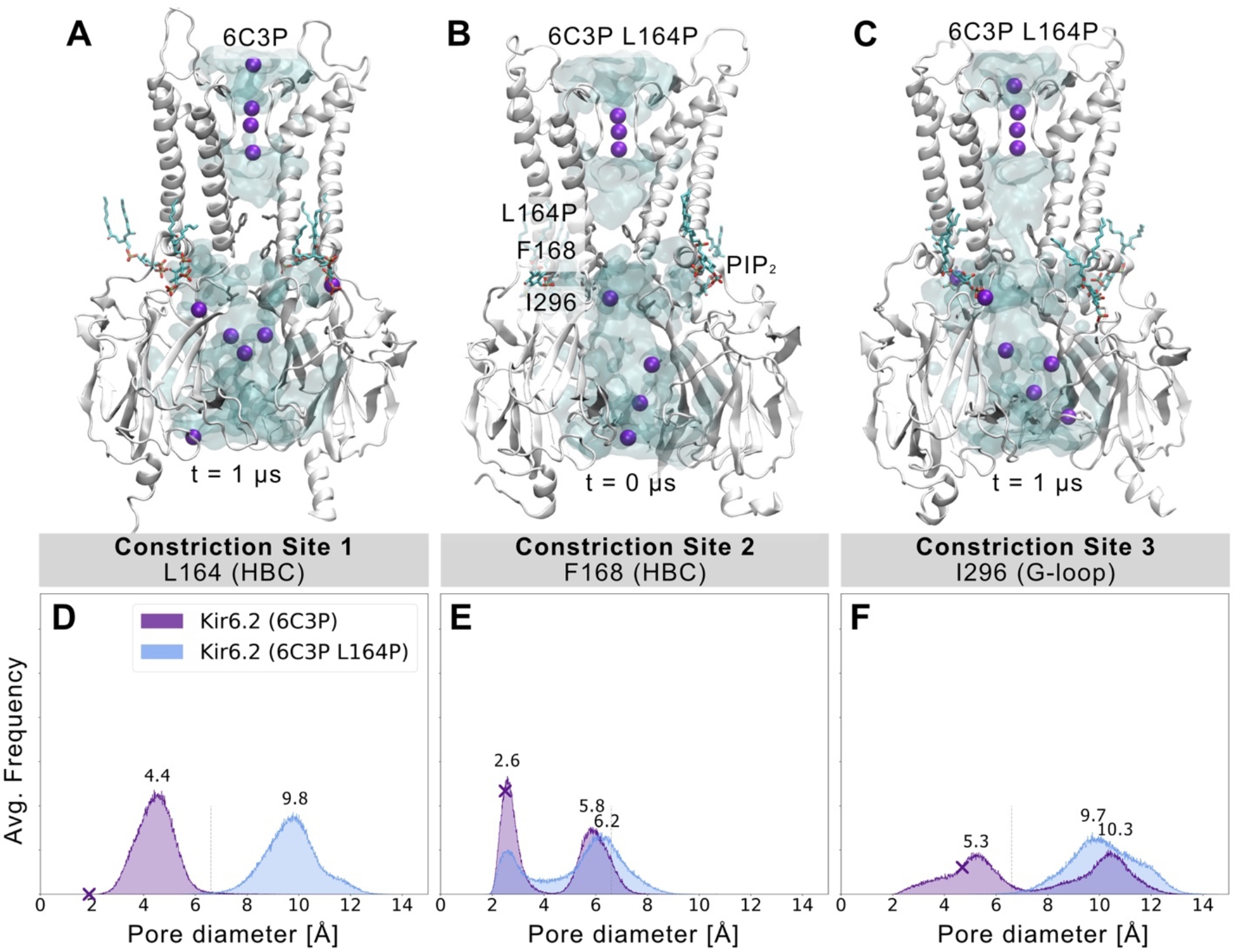
L164P leads to widening of the gates and pore solvation. Snapshots of (A) a WT 6C3P system and the equilibrated mutant system 6C3P L164P (B) before and (C) after 1 μs simulation show the solvation state of the HBC gate. The G-loop gate stabilized fully in an open conformation. (D-F) Like described in Figure 2, histograms of pore diameters show minimum distances for the three major constriction sites of 6C3P and 6C3P L164P. Constriction site 1 and 3 sampled distances wider than the time-averaged hydration diameter of K^+^ in the mutant system. The bimodal distribution at constriction site 2 shifts towards wider apertures, leading to intermittent solvation of the HBC gate. Distances were measured in 2 x 1 μs MD simulation for both system between two opposing subunits and averaged over the number of simulations, subsequently.

Since proline is known to induce kinks in helices (Cordes, Bright, and Sansom 2002), we analyzed the maximum helix angle along the length of the M2 helix over time. Supplementary Figure 5 shows an increase in the maximum M2 helix angle from ~10° to more than 15° on average between WT (6C3O and 6C3P) and L164P mutant simulations. This kinking of M2 helices and the lack of the hydrophobic side chain at constriction site 1 lead to a widening of the pore. Interestingly, previous studies reported that mutating L164 to cysteine, alanine, valine, threonine or glycine also leads to a large increase in Po (Enkvetchakul et al. 2000, 2001; Loussouarn et al. 2000, 2001) and can reduce block by verapamil (L164C)(Ninomiya et al. 2003) and chloroquine (L164A)(Ponce-Balbuena et al. 2012). Moreover, patients carrying the L164P mutation could not convert to sulfonylurea therapy. The fact that mutant channels are insensitive to regulation by these drugs, SUR, and ATP regulation (Tammaro et al. 2008) corroborates the relevance of L164 in Kir6.2 gating. Our simulations suggest a link between increased open probability and changed pore geometry. Nevertheless, future studies, including docking and drug screening, will be necessary to address this issue in more detail.

## Conclusion

Our study provides mechanistic insights into how PIP_2_ influences the narrow constrictions sites observed in Kir6.2 cryo-EM structures and unravels first structural changes induced by the permanent neonatal diabetes mutation L164P on an atomistic level.

## Materials and Methods

### Pore analysis with HOLE

Available molecular assemblies of Kir6.2 from the PDB (Research Collaboratory for Structural Bioinformatics Protein Data Bank (RCSB PDB), RRID:SCR_012820) (see Table 1), were aligned with the Swiss-PdbViewer (Swiss-PdbViewerDeepViewv4.0, RRID:SCR_013295)(Guex and Peitsch 1997) at the selectivity filter motif TTIGYG. Hydrogens were added to the pore-only files with the help of the APBS-PDB2PQR web server (Adaptive Poisson-Boltzmann Solver, RRID:SCR_008387)(Baker et al. 2001; Dolinsky et al. 2004) with standard settings (pH 7.0, PARSE FF, internal naming scheme, neutral N and C termini). Slices through the pore forming Kir channel surfaces were generated with PyMOL (PyMOL, RRID:SCR_000305)(Schrödinger 2015). The surface is colored according to the APBS map with grid spacing of 0.5. The pqr files were used to calculate the pore dimensions of the Kir channels with the HOLE program (version 2.0) (Smart et al. 1996). Standard settings were used, with pore ending radii (endrad) of 6 Å in channels. The HOLE pore geometry was analyzed with Matplotlib (MatPlotLib, RRID:SCR_008624)(Hunter 2007) and visualized with VMD (Visual Molecular Dynamics, RRID:SCR_001820)(Humphrey, Dalke, and Schulten 1996).

### MD simulations

Four short-chain C8-PIP_2_ molecules were inserted in both Kir6.2 systems (PDB accession no. 6C3O and 6C3P) based on the respective binding conformations, using the 3SYA structure (Whorton and MacKinnon 2011) as a template. PIP_2_ parameters were taken from our previous work (S. J. Lee et al. 2016). For our MD simulations, we used the amber99sb force field (Hornak et al. 2006) with Berger lipid parameters (Berger, Edholm, and Jahnig 1997). Further parameters can be found in our previous work (Bernsteiner et al. 2019). Protein topologies were created with the Gromacs module pdb2gmx (Abraham et al. 2018). The PIP_2_-bound or apo 6C3O structure was placed in the pre-equilibrated POPC (1-palmitoyl-2-oleoyl-sn-glycero-3-phosphocholine) membrane of our Kir3.2 system (Bernsteiner et al. 2019), containing 588 lipids. K^+^ ions were placed at selectivity filter positions (0,2,4), separated by water molecules. After solvation with 82,131 water molecules (SPC/E) (Berendsen, Grigera, and Straatsma 1987) and neutralization with K^+^ ions, 150 mM KCl was added to the simulation system. The energy was minimized (steepest descent) in all setups, followed by 1 ns NVT and at least 10 ns NPT equilibration until convergence of temperature and pressure, with restraints on protein atoms to their starting positions (force constant of 1,000 kJ mol^-1^ nm^-2^). 6C3P systems were set up in the same way, while the L164P mutation was introduced into Kir6.2 with the Swiss PdbViewer (Guex and Peitsch 1997). We chose 6C3P for the mutation, as the model exhibited a wider pore geometry at the cryo-EM state. Gromacs version 2018.9 (GROMACS, RRID:SCR_014565)(Abraham et al. 2015, 2018) was used to perform all-atomistic MD simulations (see table 2 for an overview). In a single 6C3P L164P simulation an electric field of 40 mV nm^-1^ along the z-axis of the simulation box was applied to test K^+^ conductivity. With a simulation system size of ~17.3 nm in the z-direction, this resulted in an electric transmembrane potential of ~700 mV (Roux 2008; Treptow and Tarek 2006). Figures were rendered with VMD (Humphrey et al. 1996) and PyMol (Humphrey et al. 1996) and movies were produced using Molywood (Wieczór et al. 2020). Maximum helix angles were calculated with the VMD Bendix plugin (Dahl, Chavent, and Sansom 2012). All data were plotted with Matplotlib (Hunter 2007).

### Minimum distance analysis

Minimum distances between opposing subunits of pore-forming Kir subunits were calculated with the Gromacs module gmx mindist (Abraham et al. 2018) for residues L164, F168 and I296 in Kir6.2, and corresponding residues in Kir3.2 (V188, F192, M319). The minimum distances of all simulations of a simulation system and both opposing subunits were combined and averaged over the number of simulations.

### PIP_2_ hydrogen bond analysis

We screened the MD trajectories for residues that formed frequent h-bonds (cut-off angle 30°, cut-off radius: 0.35 nm) between PIP_2_ and Kir6.2 with the VMD Hydrogen Bonds extension. Selected h-bonds were further analyzed with Gromacs, using the hbond module.

### Relative TMD-CTD rotation

The relative rotation of a single subunit’s cytoplasmic domain (CTD) with respect to its transmembrane domain (TMD) was calculated with an in-house script, as described previously (Bernsteiner et al. 2019). The torsional angle was defined by four points: The center of mass (COM) of the TMD’s subunit 1 (point 1), the COM of the whole TMD (point 2), the COM of the whole CTD (point 3) and the COM of CTD’s subunit 1 (point 4). Kir6.2 residues 53-98 and 116-172 were defined as TMD, while residues 32-52 and 173-346 contributed to the CTD. Thus, the highly flexible extracellular loops in the TMD and the C-termini were excluded for the COM calculations (see supplementary Figure 4E).

## Acknowledgments

The computational results presented have been achieved in part using the Vienna Scientific Cluster (VSC).

## Conflict of Interest

The authors declare that the research was conducted in the absence of any commercial or financial relationships that could be construed as a potential conflict of interest.

## Author Contributions

AS-W designed the study. MB and SP performed simulations. MB, SP and AS-W analyzed the data. MB and AS-W wrote the paper.

## Funding

This work was supported by the doctoral program “Molecular drug targets” W1232, from the Austrian Science Fund (FWF; http://www.fwf.ac.at).

## Notes

### Competing Interest Statement

The authors have declared no competing interest.

