## Supplementary Material for "Simulating PIP_2_-induced gating transitions in Kir6.2 channels"

### 1 Supplementary Figures

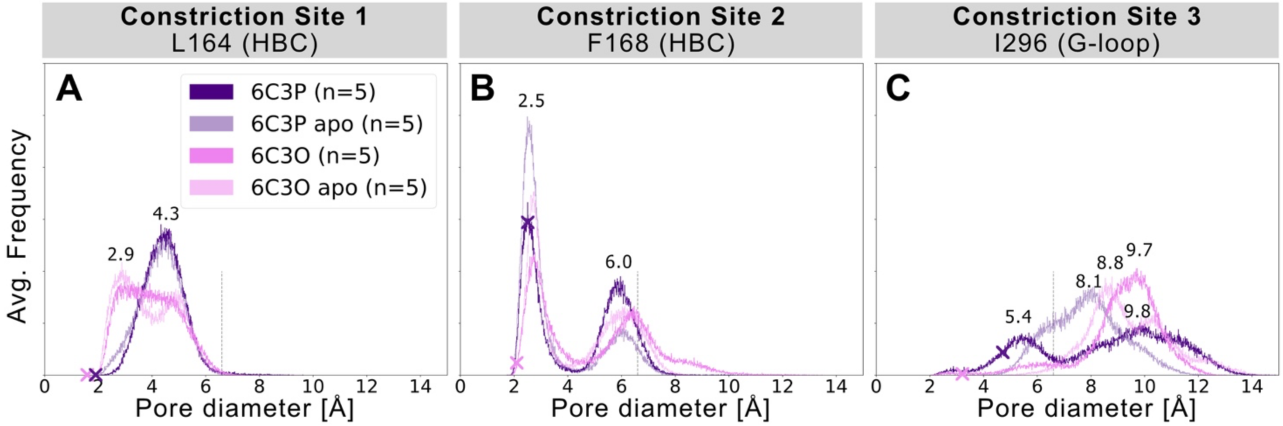


**Supplementary Figure 1.** ***Comparison of gate minimum distances in $PIP_2$-holo and apo systems.*** Minimum distances for three major constriction sites in Kir6.2 were measured in 5 x 200 ns MD simulation for all systems between two opposing subunits and subsequently averaged over the number of simulations. Like shown in Figure 2, crosses mark the corresponding distance in the initial state of the cryo-EM structures before equilibration and production run, measured with the HOLE program. A vertical line is drawn at 6.6 Å, indicating the time-averaged hydration diameter of K+ (Conway, 1981).

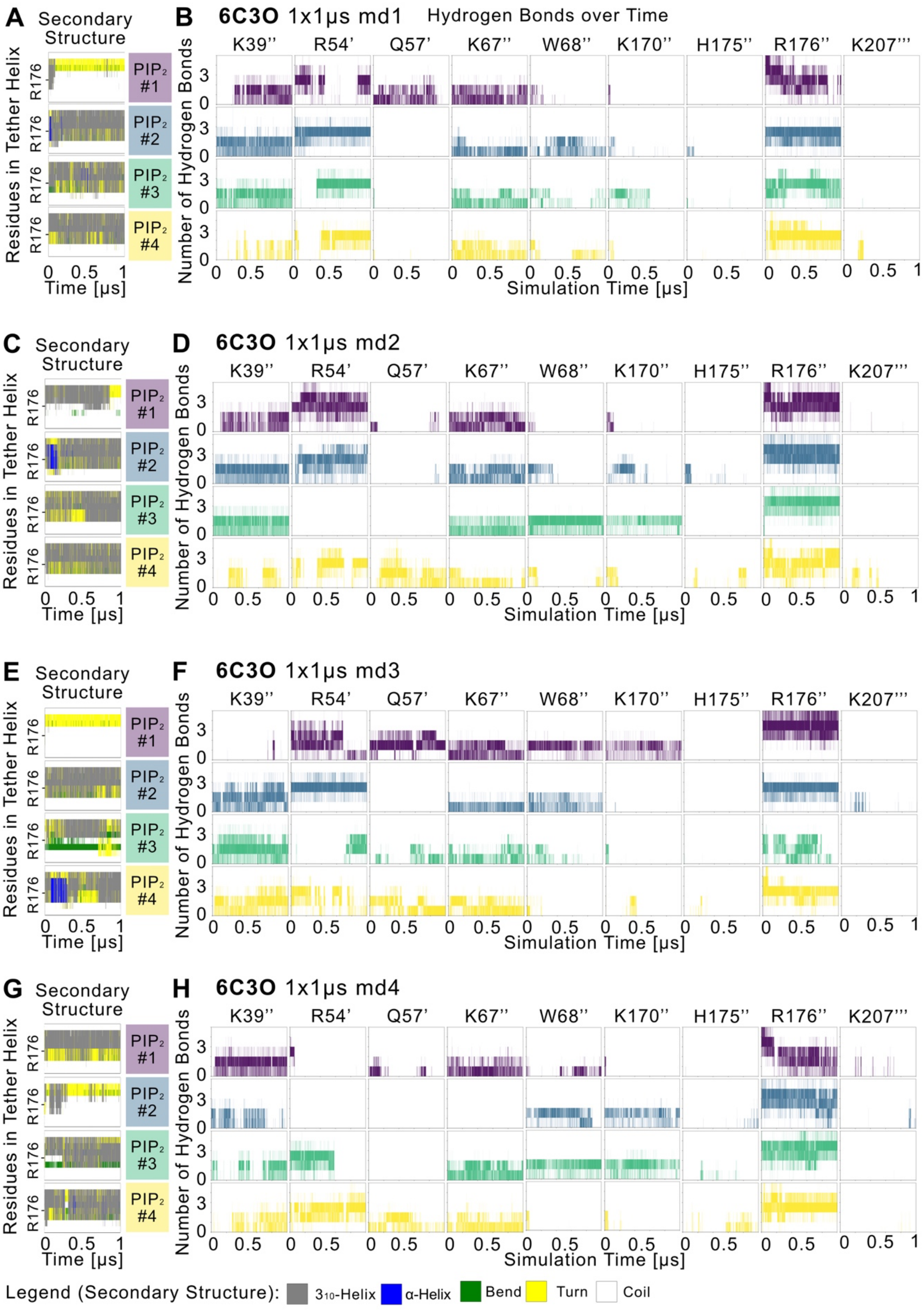


**Supplementary Figure 2.** ***PIP$_2$ binding site and PIP$_2$-induced gating changes for the remaining 6C3O simulations.*** For a description, please see Figure 3.

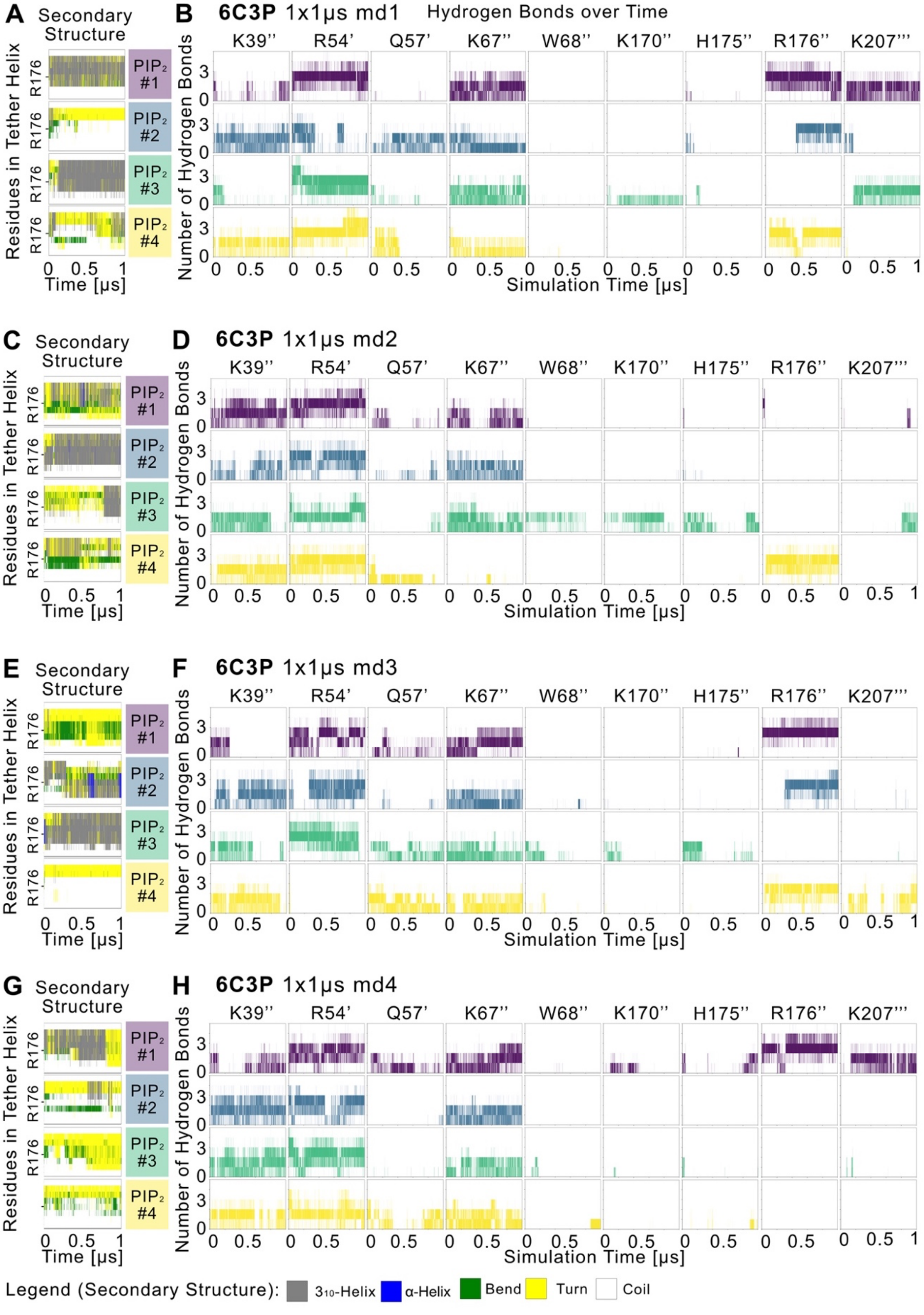


**Supplementary Figure 3.** ***$PIP_2$ binding site and $PIP_2$-induced gating changes for the remaining 6C3P simulations.*** For a description, please see Figure 3.

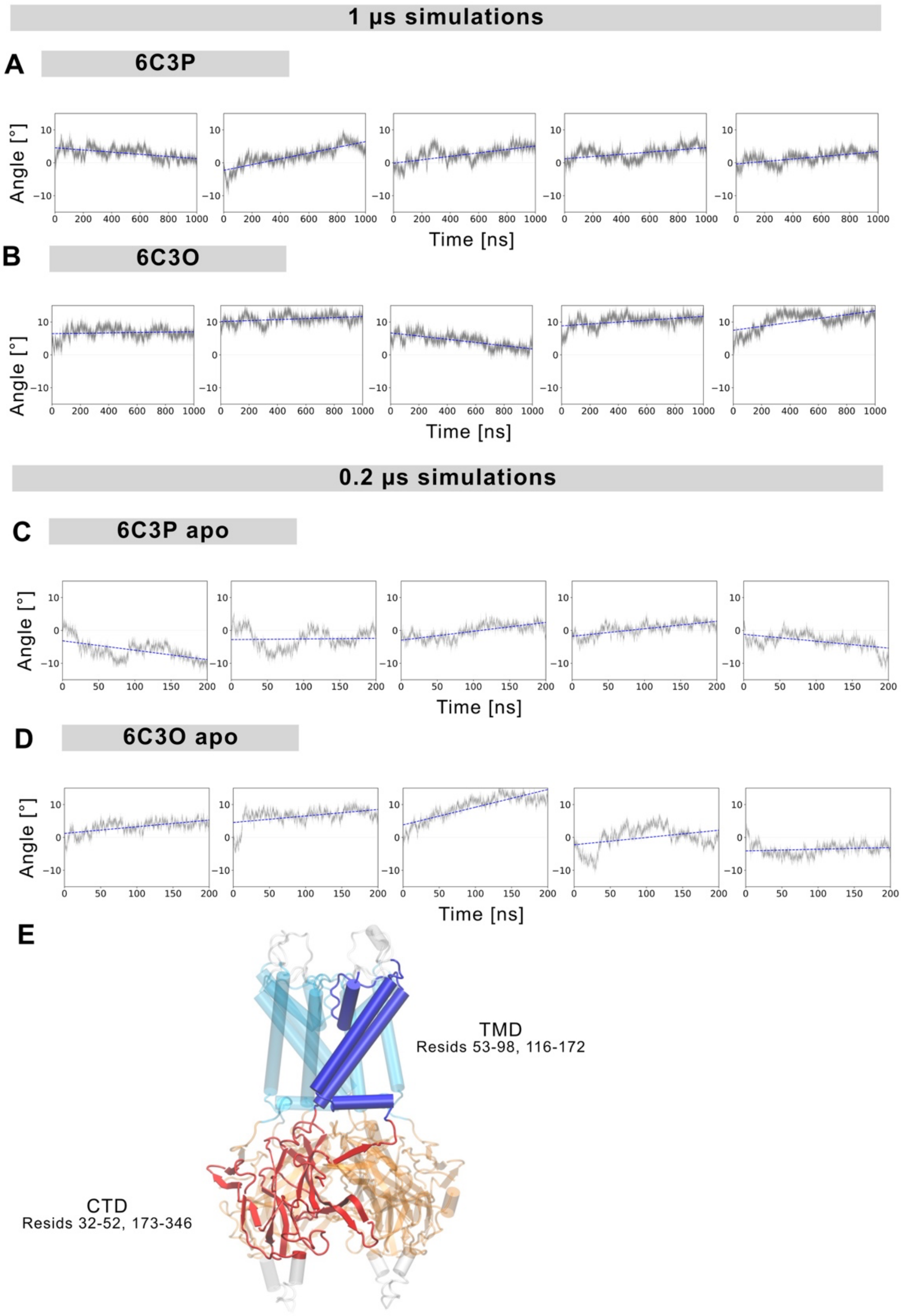


**Supplementary Figure 4.** ***Relative TMD-CTD rotations.*** Relative rotation angles as a function of time are shown for (A,B) 1 μs and (C,D) 0.2 μs simulations of different systems. The absolute angles were subtracted from the initial angles (6C3O: 60.76°, 6C3P: 68.24°), measured before the simulations. (E) Schematic figure illustrating the parts of the protein that were used to calculate the center of mass (COM) for the TMD-CTD rotation, as described in Materials and methods.

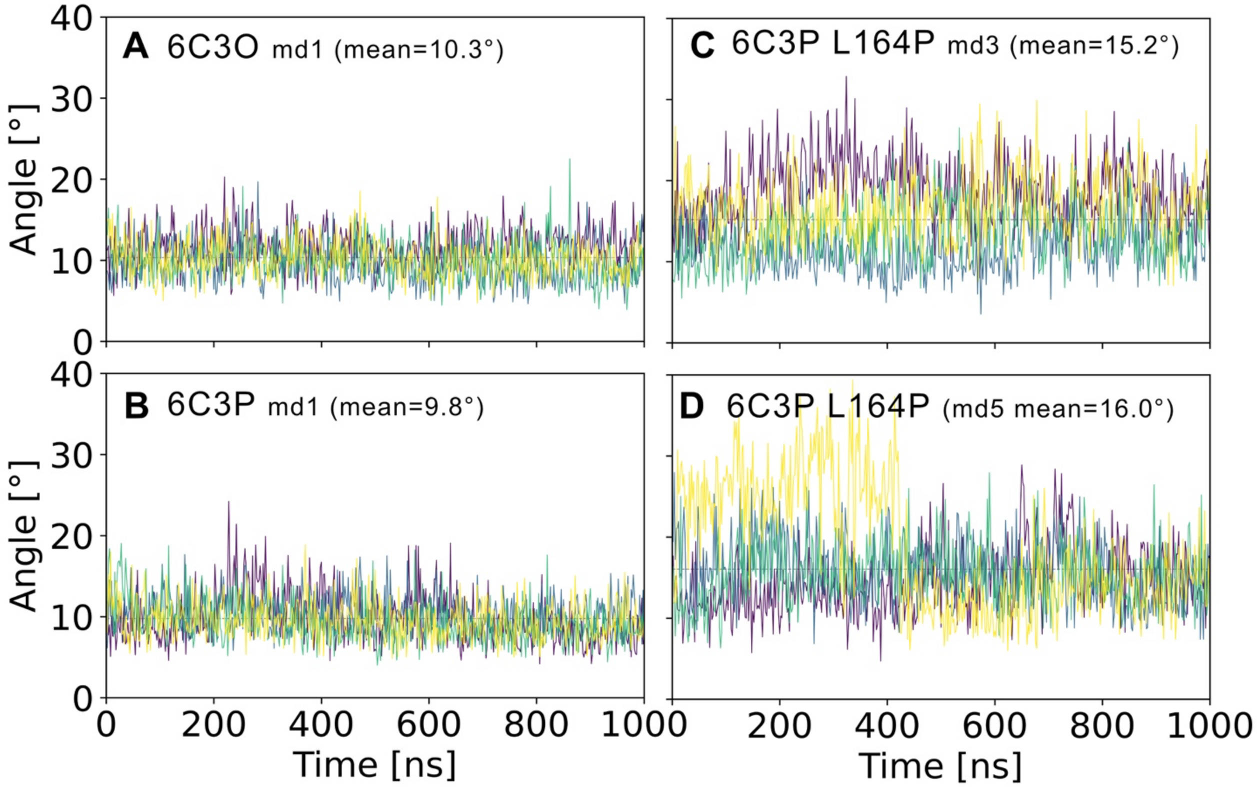


**Supplementary Figure 5.** ***L164P induces a kink in the M2 helix.*** The maximum angle of the M2 helix (residues 143-172) were calculated with the Bendix plugin of VMD (Dahl et al., 2012) for (A) a 6C3O (B) a 6C3P (C,D) two 6C3P L164P systems over time. The colors represent different subunits of the protein.

### 2 Supplementary Movies

**Supplementary Movie 1.** ***Gating and pore solvation in 6C3O***

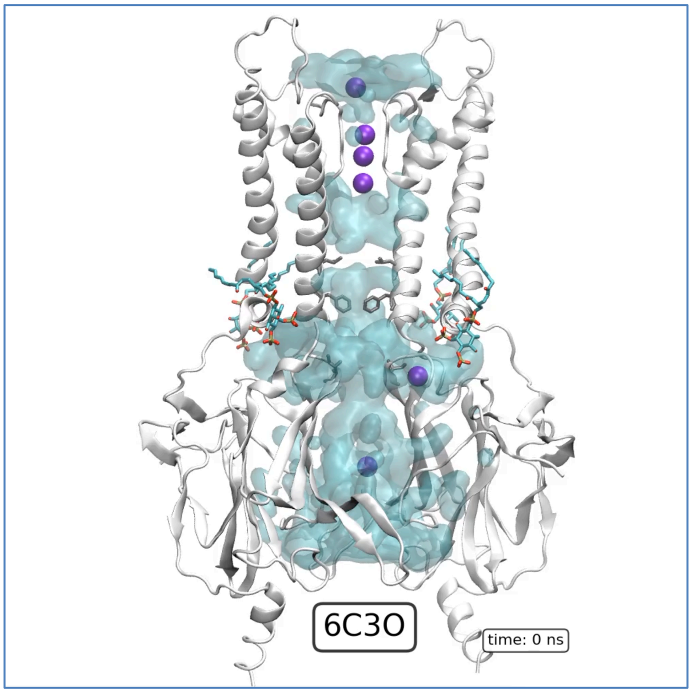


**Supplementary Movie 2.** ***Gating and pore solvation in 6C3P***

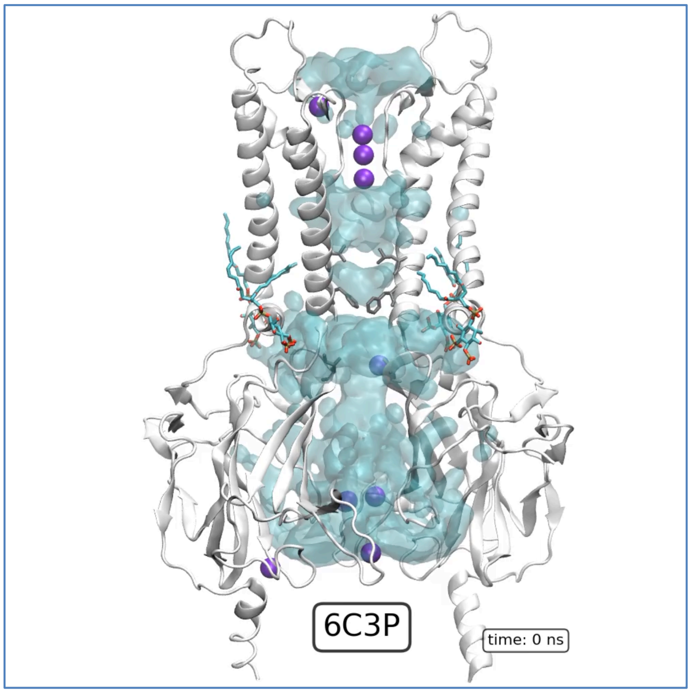

**Supplementary Movie 3.** ***Gating and pore solvation in 6C3P L164P***

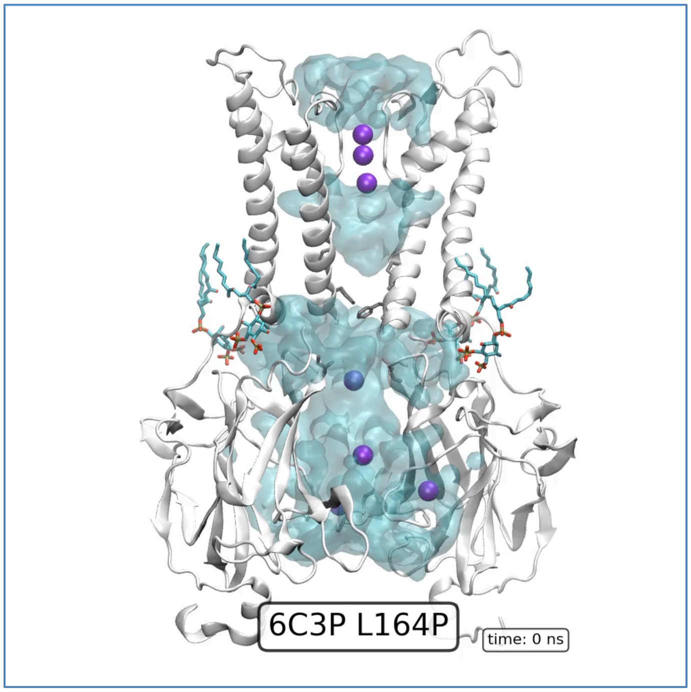
